# Spatial sampling of deep neural network features improves encoding models of foveal and peripheral visual processing in humans

**DOI:** 10.1101/2024.08.05.606515

**Authors:** Niklas Müller, H. Steven Scholte, Iris I. A. Groen

**Affiliations:** Psychology Research Institute, University of Amsterdam, The Netherlands; Informatics Institute, University of Amsterdam, The Netherlands Psychology Research Institute, University of Amsterdam, The Netherlands

## Abstract

Deep Neural Networks (DNNs) are increasingly being used to build encoding models to predict neural data recorded during visual stimulation. Their ability to process natural images makes them a prime candidate for studying the neural profile underlying real-world visual perception. However, there are still prominent discrepancies in how DNNs and humans process visual information. One discrepancy lies in the spatial sampling of visual input: while DNNs uniformly sample from their input at every spatial location, the human visual system samples differentially from central and peripheral regions. Here, we implement multiple spatial sampling strategies on feature maps of DNNs into encoding models that predict human EEG responses to a novel stimulus set consisting of large, high-quality natural scene images. By applying differential spatial sampling of DNN feature maps, we reveal distinct temporal profiles for encoding of peripheral vs. central information in EEG signals. Moreover, we show that a differential spatial sampling derived from the density of retinal ganglion cells yields the best performing encoding model when using DNN feature maps. We experimentally confirm this pattern by separately stimulating peripheral and central visual field regions, and demonstrate that the distinct temporal profiles for central and peripheral information are only revealed when using large-field stimuli. Together, these results show that aligning the spatial sampling of humans and DNN encoding models can improve predictions of neural data. The distinct temporal profiles for encoding of peripheral vs. central information support a global-to-local processing hierarchy of real-world vision.

## 1 Introduction

Deep Neural Networks (DNNs) can be used to make predictions of neural recordings during visual stimulation with high accuracy (van Gerven, 2017). Encoding models built on DNN feature maps allow for image-computable predictions of neural data across multiple species and modalities - including monkey electrophysiology (Yamins et al., 2014; Cadieu et al., 2014), human MEG (Seeliger et al., 2018), BOLD (Gü çlü and Van Gerven, 2015; Storrs et al., 2021), and EEG (Gifford et al., 2022). These models can be used to test hypotheses by changing specific aspects about the DNN architecture, training data or training regime, and subsequently evaluating whether these increase representational alignment with human behavior or neural responses (Sucholutsky et al., 2023). While such tests have inspired a range of mechanistic models for phenomena of visual processing (Doerig et al., 2023), the main contributor to the representational alignment between DNNs and humans appears to be more the high dimensional nature of deep hierarchical convolutions (even for randomly initialized weights) and less the task-optimization of convolutional features (Kazemian et al., 2024; Elmoznino and Bonner, 2024) or the DNN architecture or training regime (Conwell et al., 2022). Here, we explore another avenue towards increasing alignment: extending neural encoding models by more explicitly implementing knowledge about human visual pathways.

One particular structural property of human visual processing is the distinction between a parvocellular and a magnocellular pathway (Rodieck, 1979). The parvocellular pathway is characterized by processing inputs with high-acuity from foveal regions while the magnocellular pathway processes inputs with low-acuity from peripheral regions. The density of retinal ganglion cells (RGCs) matches the information density for foveal and peripheral inputs, yielding an overrepresentation of the fovea in the RGC output compared to the periphery (Oyster et al., 1981; Wässle et al., 1989; Kwon and Liu, 2019). The abundance of foveally sampled information is in turn compensated for by assigning substantially more brain volume to the processing of foveal information compared to peripheral information (cortical magnification, Cowey and Rolls, 1974). Previous work has shown that leveraging this property by training DNNs on foveated stimuli increases robustness to occlusions during scene categorization tasks (Deza and Konkle, 2020).

Despite the foveal overrepresentation, peripheral information serves important roles in visual perception and navigation (Holmes et al., 1977; Schyns and Oliva, 1994; Rosenholtz, 2016; Loschky et al., 2017; Vater et al., 2022). The fast magnocellular pathway (Delorme et al., 1999) aids the processing of scene gist (Potter, 1976; Biederman, 1981; Trouilloud et al., 2020) and motion detection (Burr and Ross, 1986). Training DNNs on blurred images (as a proxy for the peripheral percept) has not only been shown to increase robustness to noise during object recognition, but also shape-bias and predictivity of human fMRI and monkey electrophysiology data (Jang and Tong, 2024). Recent work from da Costa et al. (2024) integrated foveal and peripheral sampling using spatial reweighting into a DNN, leading to the emergence of a hierarchy of receptive field properties similar to that of the human visual system. However, the implications of cortical magnification on predicting neural responses have so far not been systematically evaluated.

Here, we aimed to explicitly test whether incorporating human-like spatial-sampling into DNN-based encoding models increases representational alignment. Using electroencephalography (EEG) recordings from human participants performing a rapid-serial-visual-presentation task (RSVP, Intraub, 1981) with natural scene images from a custom dataset of large, high quality photographs, we show that spatial feature selection significantly improves encoding models based on DNN features. We find differential contributions of peripheral vs. central information in the scene to the encoding performance of EEG recordings over time, consistent with theories of coarse to fine visual processing in which a global precedes a local, detailed percept (Bar, 2004; Hegdé, 2008). Importantly, we confirm our modeling results with an additional experiment that selectively stimulates foveal and peripheral regions, and show that using large-field visual stimulation, more closely simulating real-world visual scenarios, is critical to reveal these temporally distinct encoding profiles.

In sum, by comparing encoding models with different spatial sampling strategies, we find clear pointers that spatially distinct regions of the visual field elicit temporally distinct neural signatures. Using a spatial transform that is based on empirical data of the distribution of RGCs in the human retina (da Costa et al., 2024), we further improve encoding performance while unifying both temporally specific dynamics.

## 2 Methods

### 2.1 Data Collection

#### 2.1.1 Main Experiment

##### Subjects

Human EEG data was collected from 31 participants (18-30 years) with normal or corrected-to-normal vision. The Ethics Review Board of the University of Amsterdam approved the experiment and all participants gave written informed consent before participation and were rewarded with study credits. Four subjects were excluded from analysis because their recordings were incomplete.

##### Stimulus set

The Open Amsterdam Data Set (OADS), a new stimulus set consisting of 6130 ultra-high resolution outdoor scenes was used for both human data collection and DNN feature extraction. OADS images consist of photographs of diverse street scenes taken in the city of Amsterdam, The Netherlands, have an original resolution of 5468×3672 pixels, and are stored in an uncompressed format. Additionally, a set of 27 indoor scene images with the same resolution depicting office scenes served as target images in the EEG experiment.

##### Experimental Procedure

Subjects were presented with scene stimuli downsampled to a resolution of 2155×1440 pixels using a rapid serial visual presentation (RSVP) paradigm (Intraub, 1981; Keysers et al., 2001; Grootswagers et al., 2019). Stimuli were shown on a 24-inch monitor with a resolution of 2560×1440 pixels and a refresh rate of 144 Hz. Subjects were seated 63.5 cm from the monitor and were using a head rest such that stimuli spanned 50×29.5 degrees of visual angle. In total, the stimulus set used in this experiment consisted of 4800 images of which every subject was presented with a subset of 702 images. 1/3 of all images were repeated 10 times throughout the entire experiment while the other 2/3 were repeated 5 times. The resulting EEG data were averaged across repetitions for each unique image to increase the Signal-To-Noise-Ratio (SNR). Following standards in the linearized encoding literature (e.g., Gifford et al., 2022), we created a low-SNR train image set to fit the encoding model by averaging across the 5 repetitions stimuli and a high-SNR test image set by averaging across the 10 repetitions stimuli.

Each subject performed 240 trials. In every trial, a series of 20 images of either only outdoor scenes (non-target trial) or 19 outdoor scenes and one indoor scene (target trial) was presented, with half of the trials being target-trials. Participants were asked to perform a target detection task by pressing either ’J’ for a target trial or ’K’ for a non-target trial, at the end of the trial. In each trial, subjects were first presented with 3500 ms gray screen and a red fixation cross in the center of the screen which was continuously presented during the entire trial. Next, 20 stimulus images were presented for 100 ms and a 300 ms gray screen each. At the end of the trial, subjects had 3500 ms to respond with a button press and were instructed to blink before the start of the next trial. The session was divided into 8 blocks each followed by an enforced break of at least 30 seconds with self-paced continuation. Stimuli were presented and behavioural data collected using *PsychoPy* (Peirce et al., 2019).

##### EEG Acquisition and Preprocessing

EEG data was collected using a Biosemi 64-channel ActiveTwo EEG system (Biosemi Instrumentation) with an extended 10–20 layout, modified with two additional occipital electrodes (I1 and I2, while removing electrodes F5 and F6). Eye movements were monitored with electro-oculograms (EOGs). EEG recording was followed by offline re-referencing to external electrodes placed on the earlobes. Resulting EEG data was pre-processed, according to the lab’s standard pipeline (e.g., Groen et al., 2012) using the MNE software package (Gramfort et al., 2013), as follows: high-pass filter at 0.1 Hz (12 dB/octave) and a low-pass filter at 30.0 Hz (24 dB/octave) followed by two notch filters at 50 and 60 Hz; automatic removal of deflections > 300*mV* ; epoch segmentation in -100 to 400 ms from stimulus onset; occular correction using the EOG electrodes (Gratton et al., 1983); baseline correction between -100 and 0 ms; and conversion to Current Source Density (Perrin et al., 1989). As explained above, the resulting ERP data were averaged across repetitions per unique image, thus resulting in an sERP specific to each subject, electrode and image.

#### 2.1.2 Additional Experiments

##### Subjects

Human EEG data was collected from 4 new participants (2 females, age range = 21-30 years) in 6 sessions. The institution’s Ethical Committee approved the experiment and all participants gave written informed consent before participation and were rewarded with 180 .

##### Stimulus set

We used a random subset of 697 outdoor scenes from the same stimulus set described in 2.1.1. Additionally, a set of 35 indoor scene images depicting office scenes served as target images in the additional experiments.

##### Additional Experiment 1 - Central and Peripheral Stimulation

For each stimulus image, 3×2 additional versions were made. Using a circular aperture of increasing diameter (3^◦^, 11^◦^, and 20^◦^), we created a center crop version that removed everything outside of the circular aperture (thus making it the same gray as the screen background) and a periphery crop version that removed everything inside the circular aperture. For each image and each of the three aperture sizes there are thus 2 versions, one containing only the parts of the image around the center, and one containing only the parts of the image in the periphery, but not in the center.

##### Additional Experiment 2 - Stimulus Size

Additionally to the periphery/center crop stimuli outlined above, we created 2 more versions for each stimulus image, changing the image size from the original 2155×1440 pixels (large) to 1079×720 pixels (medium), and 529×360 pixels (small). All three size condition were included in the experiment. Thus, for each original image, 9 different stimulus conditions were presented.

##### Experimental Procedure

This experiment was conducted using the same experimental paradigm as the main experiment (see 2.1.1). The following details differed in the additional experiments: all 4 subject saw the same set of 6273 (697 images x 9 conditions) stimuli with randomized stimulus orders per subject and session. 35% of all images were repeated 10 times while the other 65% were repeated 5 times.

Each subject performed 2706 trials (451 per session). In every trial a series of 20 images were presented. In 50% of trials, one of the 20 images was an indoor image, in 25% of trials, two images were indoor images, and in 12.5%, three images were indoor images. Trials with one or three indoor images were defined as target trials. This changed compared to the main experiment was introduced to enhance attention, specifically because 6 sessions were performed.

##### EEG Acquisition and Preprocessing

EEG signals were recorded using the same setup as outlined in 2.1.1 with the following differences: electrodes F5 and F6 were not replaced by electrodes I1 and I2.

### 2.2 Computational Modelling

#### 2.2.1 Regression on Trial-Averaged ERPs

##### EEG encoding models

We built linearized encoding models to map convolutional features from DNNs onto the ERP amplitude for each subject, electrode and time point. All model fitting was performed using *python3.10*.

##### DNN feature extraction

We used the pre-trained features of an ImageNet-trained alexnet (Krizhevsky et al., 2012) from all max pooling layers (features.2, features.5, and features.12) using *PyTorch* (Paszke et al., 2019). Layers were selected by grouping them into ”processing block” that applied a convolution, a non-linearity and a maxpooling, of which the output was then extracted.

##### Spatial feature selection

We applied spatial feature selection on each extracted convolutional feature map. Feature selection was done in one of the following ways:

I. **center crop** - a crop of 0.5% of the area of the full 2D feature map was taken at the center of each feature map. The rest of the feature map was discarded. The absolute size of the crop depended on the layer and ranges from 18×12 pixels (layer1, 64 channels), over 9×6 (layer2, 192 channels), to 4×3 pixels (layer3, 256 channels).
II. **periphery crop** - the central 0.5% of the area of the full feature map was removed from the center of each feature map. This was thus the inverse of the center crop.
III. **Ganglion Cell Sampling** (GCS, da Costa et al., 2024) - a retinal ganglion cell transform was being applied to the full feature map. The GCS transform applies a magnification to central pixels and reduces the spatial weighting of peripheral pixels. The output of the transform is always square with the length of each side being the maximum of the input side lengths. The default parameters reported in the original paper were used with most importantly the magnification factor ”foveal size” set to 20^◦^.

As a baseline, we included the condition where the full feature map was used as input to the encoding model.

##### Model fitting procedure

The stimuli used during the EEG experiment were divided into two sets: a low-SNR set of 5 repetitions per image and a high-SNR set with 10 repetitions. Stimuli from the low-SNR set were used to fit the linearized encoding model (training set) while the stimuli from the high-SNR set were used to evaluate generalizability (test set).

For every model instance, we flattened the selected features per stimulus in the low-SNR set across all layers and performed a principal component analysis (PCA) across these features. The projections per stimulus of the first 100 PCs then formed the design matrix. The ERP amplitudes for every subject, electrode, and time point were then regressed onto the design matrix. On the selected features of the stimuli of the high-SNR test set, we applied the same PC projections and then used the fitted encoding model to predict the ERP amplitude for every subject, electrode, and time point. We then quantified each model’s predictive performance by calculating the cross-validated Pearson correlation coefficient (r) between predicted and real ERP amplitude.

##### Statistical significance

For each encoding model’s prediction performance, we performed a one-sample t-test against 0 across subjects for each channel and time point using the *scipy* software package (Virtanen et al., 2020). We applied false-discovery-rate (FDR) (Benjamini and Hochberg, 1995) correction across all test results and indicated statistical significance for *alpha* = 0.01 if not stated otherwise.

Additionally, for pairs of encoding models we compared prediction performance to determine the best performing model at each channel and time point. We performed a one-sample t-test against 0 across subjects of the differences of encoding model performances. We applied FDR-correction across all test results and indicated statistical significance for *alpha* = 0.05 if not stated otherwise.

##### Noise ceiling calculation

We calculated the noise ceiling, i.e., theoretical maximum of explainable variance of the ERP amplitude across stimuli, per subject, electrode and time point using the same method as described in Gifford et al. (2022). We split the ERP signal across ten repetitions for the high-SNR set into two non-overlapping splits of 5 repetitions each. For each subject, electrode and time point, we estimated the lower bound of the noise ceiling as the correlation between the average of the first split of 5 repetitions and the average of the second split of 5 repetitions, and the upper bound as the correlation between the average of the first split of 5 repetitions and the average of all 10 repetitions. Noise ceilings for the additional experiments per condition are omitted in Fig. 4 for clarity but can be found in Fig. S2 together with the mean ERPs per condition.

##### Partial correlation analysis

We calculated partial correlations using the *pingouin* software package (Vallat, 2018). For pairs of encoding models, we calculate partial correlations for each model’s predictions while regressing out the variance predicted by the other model, for each subject, electrode, and time point. We report partial correlation for selected channels, over time for center vs. periphery, center vs. GCS, periphery vs. GCS, and full vs. GCS model pairs.

##### Selection of electrodes and time points

The large number of data points and ensuing statistical results (31 subjects x 64 electrodes x 513 time point x 4 encoding models) required selection of representative results. For figures 2 and 3, two electrodes have been chosen to represent occipital electrodes (Iz) and more central electrodes (Pz). Additionally, for reporting topoplots in Fig. 2 we selected three time point as representatives for early (95 ms after stimulus onset), intermediate (119 ms) and late (144 ms) times. For Fig. 4 the average across the 17 posterior electrodes (O1, Oz, O2, PO7, PO3, POz, PO4, PO8, P7, P5, P3, P1, Pz, P2, P4, P6, P8) was reported (in accordance with Gifford et al., 2022). For Fig. 5 both Iz and the average across the 16 posterior electrodes were reported.

**Figure 1:**
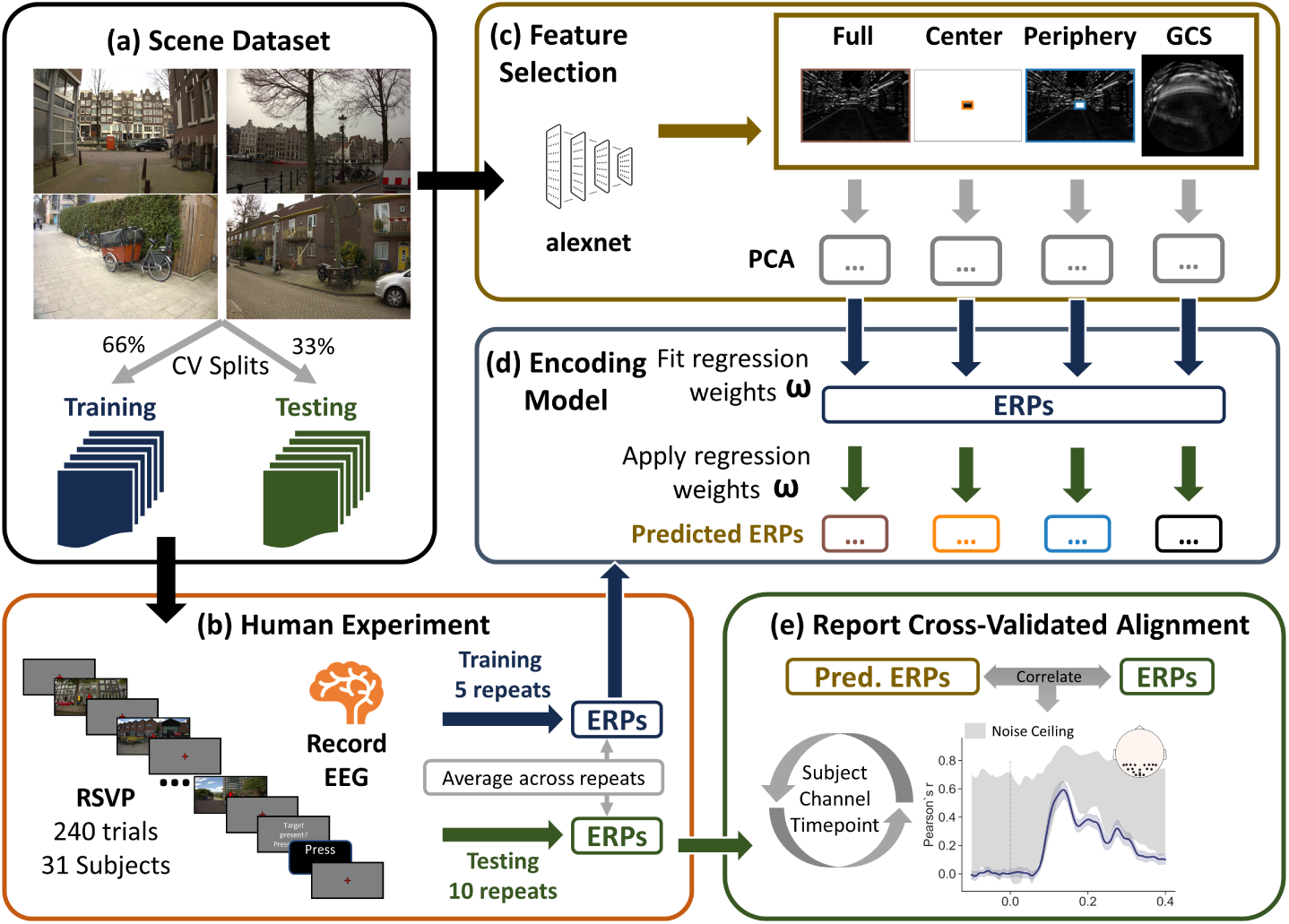
Linearized Encoding Models for Predicting Human RSVP EEG Recordings. **a)** Example stimuli used for the human EEG experiment and DNN feature extraction. The dataset was split into two non-overlapping sets: a training (66%) and a test set (33%). **b)** EEG data was recorded during an RSVP experiment with a indoor-scene detection task. **c)** Feature maps from 3 layers of a DNN were extracted per stimulus. For each feature, the full feature map was used as the baseline condition. Additionally, for each feature map, three more versions were created: (I) ”Center” - only the central 0.5% of the feature map was kept while the rest was discarded; (II) ”Periphery” - the central 0.5% of the feature map were removed while the rest was kept; (III) ”GCS” - a retinal ganglion sampling (da Costa et al., 2024) transformation was applied to the full feature map. The resulting feature maps per version were flattened followed by separate PCAs per version. **d)** Encoding models were fitted for each subject, channel, time point, and feature selection version on the training data that linearly mapped the PCA-transformed DNN features to ERP amplitudes. The fitted regression weights were then applied to the held-out features from the test set to predict ERP amplitudes for each image, per subject, channel, time point and feature selection version in (c). **e)** Predicted ERP amplitudes were compared to the recorded ERP amplitudes using Pearson correlation. The cross-validated encoding performance was reported per condition and statistical test were performed across subjects to determine the best model per time point and channel.

**Figure 2:**
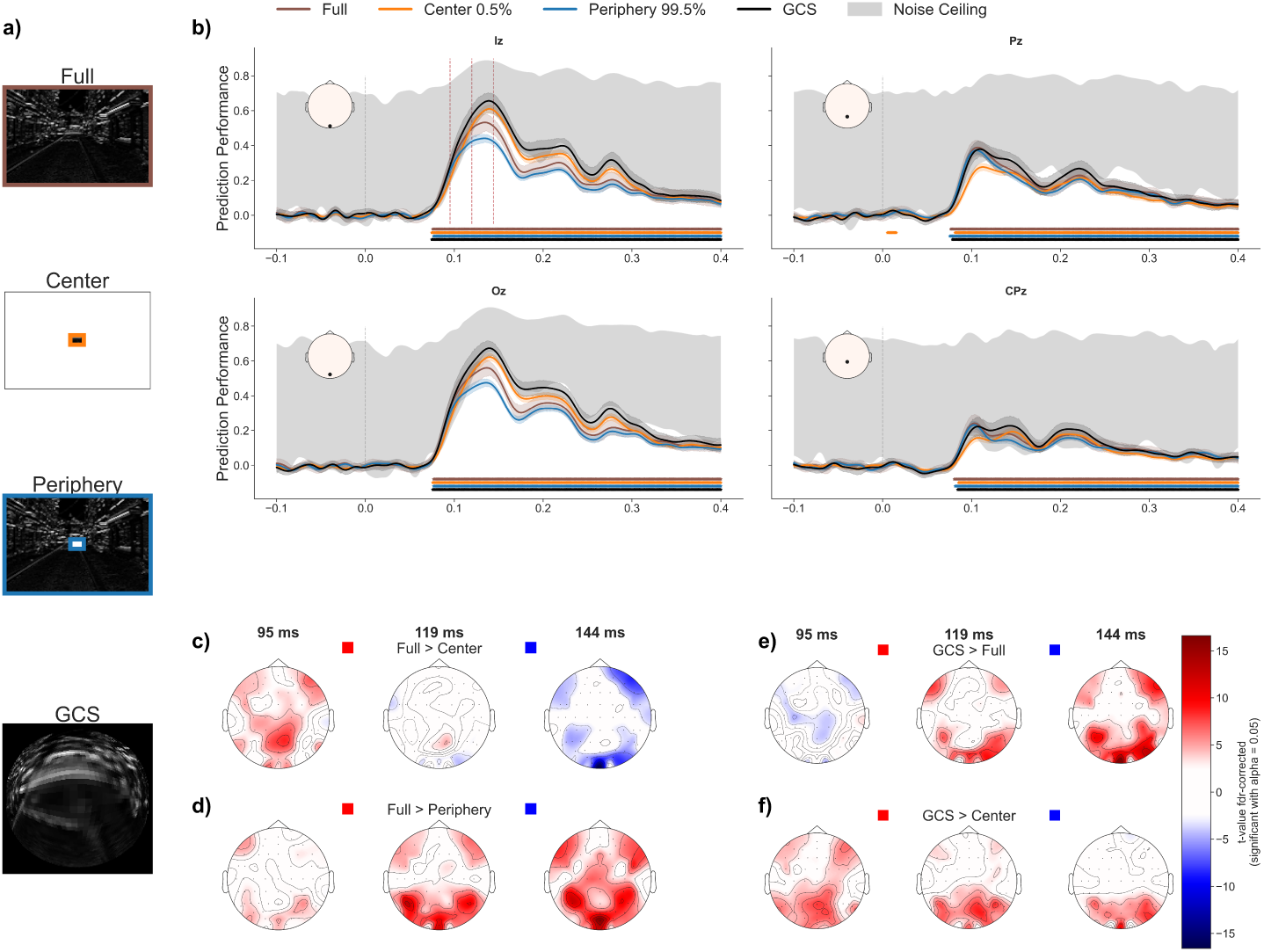
Spatial feature selection improves encoding model predictions. **a)** Examples of spatial transformations of DNN feature maps. Transforms either (I) keep the full feature map, (II) take a center-crop, (III) take a periphery-crop, i.e., removing exactly the center-crop, or (IV) apply a retinal sampling (GCS) magnifying the center while reducing, yet maintaining peripheral information (from top to bottom). **b)** Mean prediction performance (Pearson’s correlation) across subjects over time for electrodes Iz and Oz (first column) and Pz and CPz (second column) for distinct encoding models using features from each of the transformations shown in a). Shaded gray areas show the estimated noise ceilings per channel, per time point, averaged across subjects. While central information dominates at Iz and Oz and peripheral information at Pz and CPz, GCS outperforms all other transformations at all time point for the shown electrodes. **c)-f)** Topoplots showing FDR-corrected t-values of a 1-sample t-test testing for statistical difference from 0 of the difference between correlations between two conditions. Tested pairs of conditions are indicated above each row (including the color coding), for time points 95 ms (first columns), 119 ms (second columns), and 144 ms (third columns).

**Figure 3:**
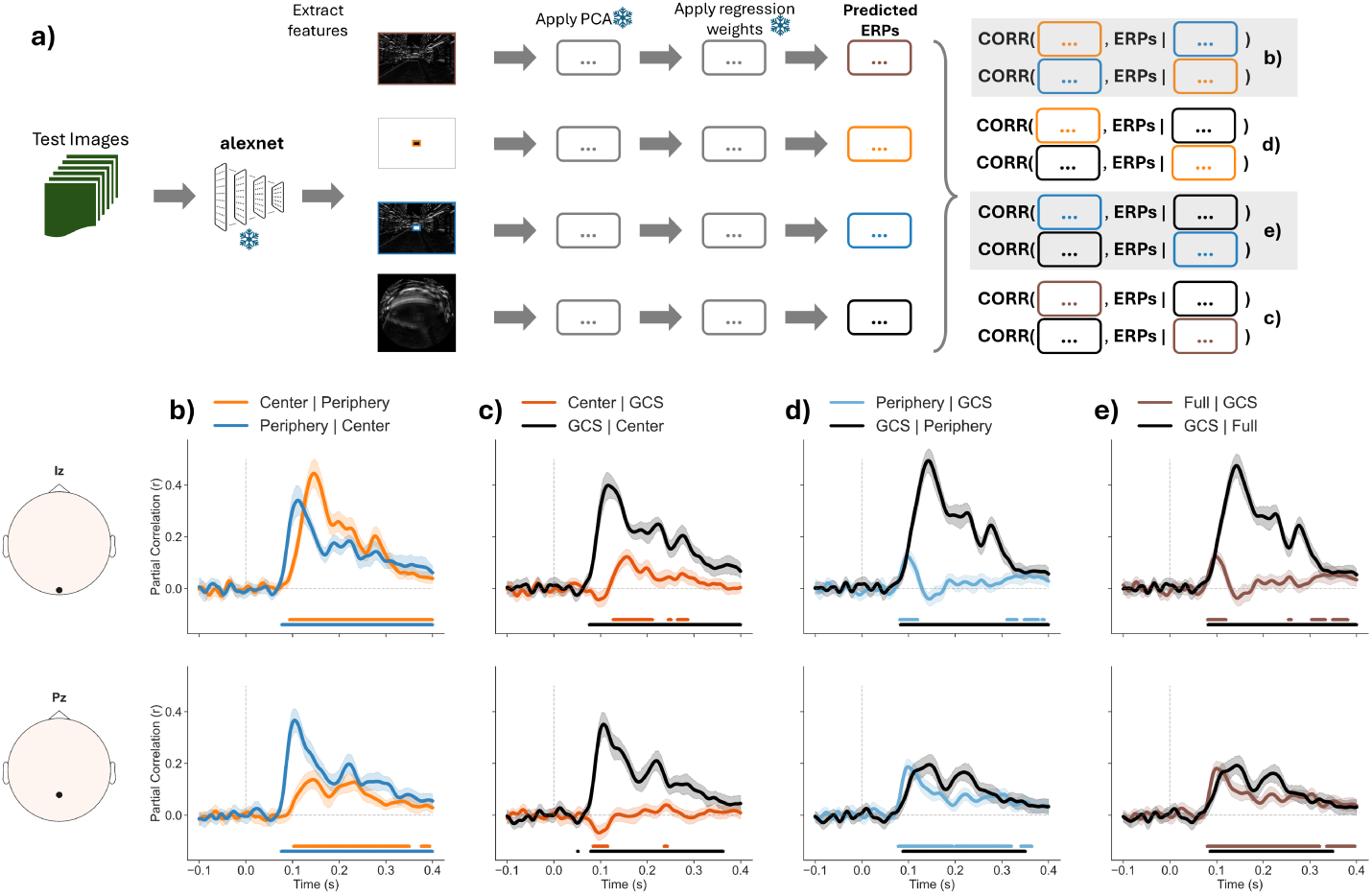
Temporally distinct ERP predictions by spatially distinct sampling, unified by retinal spatial reweighting. Plots show partial correlations (r) over time for pairs of encoding model performances. **a)** Overview of partial correlation computation. DNN features maps were extracted for test set images, spatial sampling was applied to feature maps and for each seperate sampling, and the respective PC transformations and encoding model weights that were fitted on the training data were applied to the flattened features concatenated across all layers. For four pairs of models, we calculated the partial correlation between the predicted ERPs of one model and the measured ERPs while regression out the variance explained by the predicted ERPs from the other model, in both directions, respectively. **b)** Partial correlations for center and periphery-crops: peripheral information predict early time points of ERPs while central information predicts later time points. **c)** Partial correlations for center-crop and GCS: GCS captures almost all variance that the center-crop explains. **d)** Partial correlations for periphery-crop and GCS: GCS captures almost all variance that the periphery-crop explains, but not vice-versa. **e)** Partial correlations for full model and GCS: GCS captures almost all variance that the full model explains, but not vice-versa.

**Figure 4:**
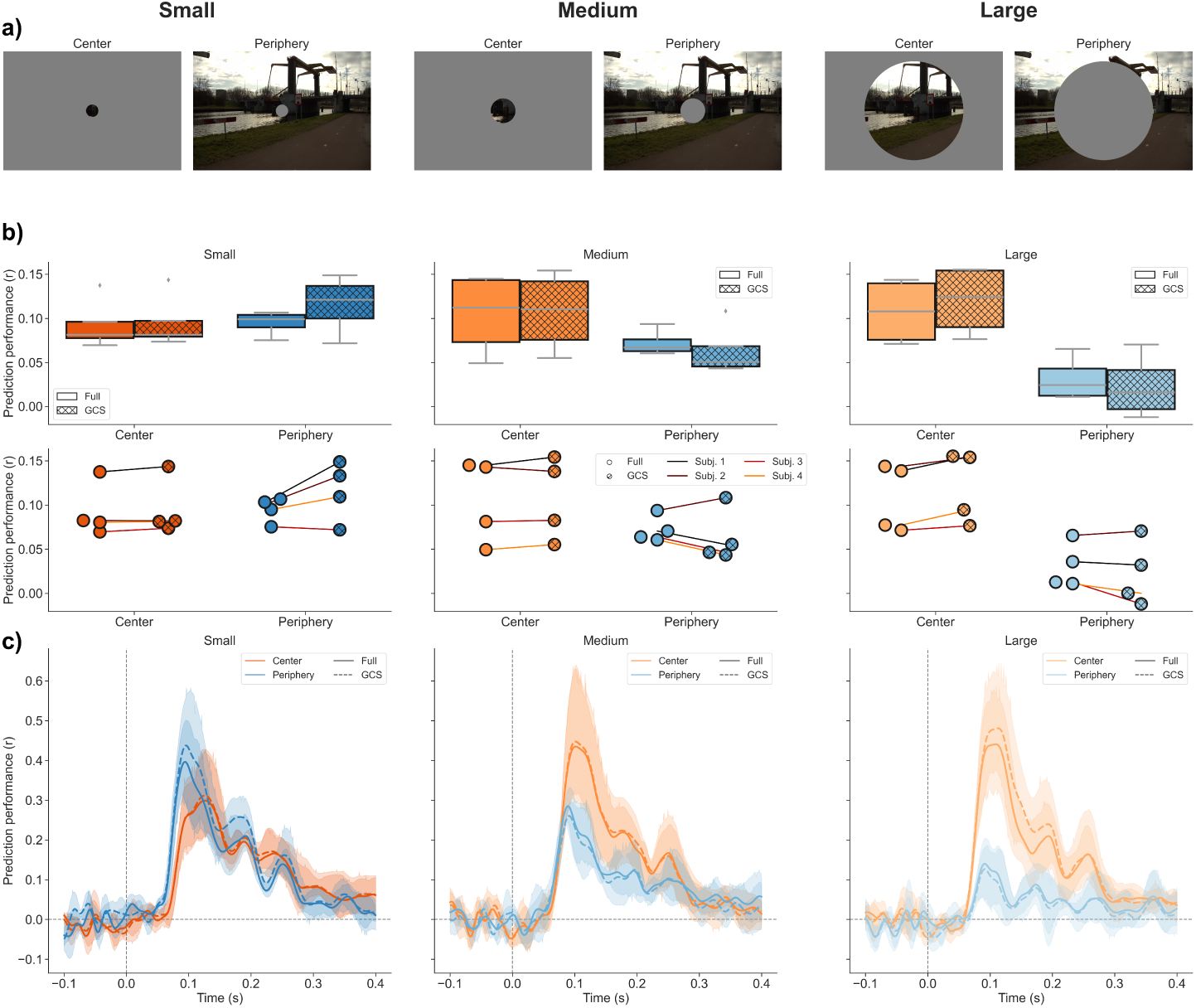
Spatially distinct stimulation elicits temporally distinct ERPs. Columns contain the example stimuli and encoding model results for center and periphery stimulus conditions using increasing circular aperture sizes: small, medium, and large (from left to right). **a)** Example stimuli for center and periphery conditions. **b)** Prediction performances across subjects for center and periphery conditions. Box and circle style indicate encoding model (”Full” vs. ”GCS”), box and circle colors indicate aperture sizes, and line colors indicate subject identity. **c)** Mean prediction performances across subjects over time. Line colors indicate aperture condition, and line style indicates which encoding model the results belong to. For small apertures, ERPs during peripheral stimulation can be predicted better, much earlier in time compared to central stimulation. GCS-transforms improves predictions most when stimulation is closest to real-world vision.

**Figure 5:**
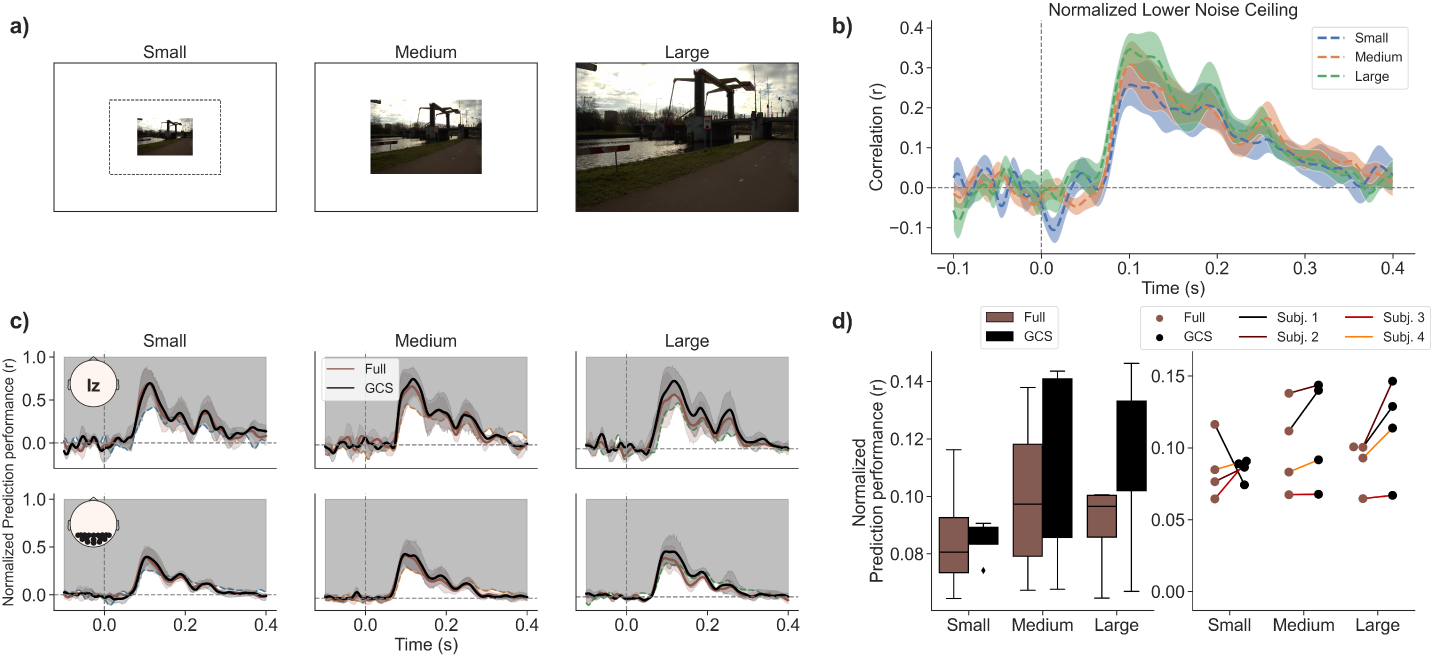
Signal-to-Noise ratio increases with size of visual stimulation. **a)** Example stimuli for the three image size conditions. **b)** Estimated lower bound of the noise ceiling normalized by the estimated upper bound averaged across 4 subjects, shaded areas show standard error, for each image size condition, respectively. **c)** Prediction performances normalized by the upper bound noise ceiling estimate over time for full-feature and GCS encoding models using ERPs from experimental runs with varying stimulus sizes. Top row shows mean performance across subject for electrode Iz. Colored shaded areas show standard error across subject per condition. Bottom row shows mean performance across subjects and the selected set of visual electrodes. Axis insets with topoplots show the locations of included electrodes on the scalp. Shaded gray areas indicated the estimated normalized noise ceiling. For each condition the normalized lower bound is colored according to b). **d)** Left panel shows boxplots of prediction performances normalized by the upper bound noise ceiling estimate for time points larger than 0 second per image size condition and feature selection version (full-feature in brown and GCS in black). Right panel shows the average performance per encoding model across channels for each subject individually, per image size condition. Dots belonging to the same subject are connected by colored lines, indicating subject identity. The noise ceiling increases with increasing stimulus size. This increased signal is captured by the GCS encoding model but not by the full-feature encoding model.

## 3 Results

### 3.1 Main Experiment

We first investigated whether selectively encoding different parts of the DNN visual feature maps improves encoding model predictions of EEG response amplitudes to natural scene images shown in an RSVP experiment (see 2.1.1). Fig. 2a) shows examples for each of the feature selections, described in more detail in 2.2.1. For each type of feature selection, we built a separate encoding model of which we show the cross-validated mean (with 95% confidence intervals) prediction performances across subjects for four selected electrodes over time in Fig. 2b).

#### 3.1.1 Spatial feature selection improves encoding model performance

We found that all model performances are significantly different from 0 for time points later than 75 ms at Iz and 76 ms at Oz (Fig. 2b, top and bottom left; insets highlight the electrode location on the scalp). For occipital electrodes Iz and Oz the cross-validated prediction performance for the center-crop model is higher than both the full model and the periphery-crop model at time points later than approximately 120 ms. The performance of the center-crop model approaches the lower bound estimate of the noise ceiling. Additionally, the performance of the periphery-crop model is lower than the full model at time points between around 120 and 145 ms.

Performances of the full and periphery model are significant at time points later than 76 ms for Pz and 80 ms for CPz (Fig. 2b, top and bottom right), while the center model is only significant after 81 ms at Pz and 85 ms at CPz. For central electrodes Pz and CPz, performance of the periphery-crop and full model are higher than the center-crop model for time points earlier than approximately 145 ms. All but the center-crop model performances seem to approach the lower bound estimate of the noise ceiling.

We statistically compare encoding model performance and thereby spatial selection methods across subjects at every electrode for the three time points 95, 119, and 144 ms after stimulus onset (time points are highlighted in line plot in Fig. 2b with vertical red dashed lines). We found that the full model significantly outperforms the center-crop model at central electrodes at 95 ms. This pattern disappears at 119 ms, when there is almost no statistically significant difference between both model’s performances. At 144 ms, the center-crop model in turn outperforms the full model at occipital electrodes and thereby confirms the intuitions observed in Fig. 2b). Thus, we find a substantial improvement of encoding model performance when selecting only the central 0.5% of the full feature map to predict human EEG data at occipital electrodes. This improvement is temporally restricted to later time points.

#### 3.1.2 Evidence for scene gist processing

Next, we compared the full model to the periphery-crop model (see Fig. 2d). At the early time point (95 ms), the full and periphery-crop model performances show limited statistically significant differences, with some occipital electrodes being better explained by the full model. At both later time points (119 and 144 ms), the full model consistently outperforms the periphery-crop model at occipital, central, and frontal electrodes. Notably, removing only 0.5% of the feature maps in the center yields significantly worse predictions compared to using the full feature map. Thus, even with 99.5% of shared information, there is a strong discrepancy between the accuracies of both encoding model predictions.

Comparing the center-crop model against the full model, as well as the periphery-crop model against the full model, allows us to compare the impact of removing either central information or peripheral information. Overall, we observe that peripheral information is sufficient and necessary to explain the majority of variance at early time points (95 ms). In turn, using peripheral information leads to poor encoding model performance at later time points while central information is now sufficient for good model performance.

#### 3.1.3 Non-uniform retinal spatial sampling holistically predicts EEG signals

So far, we have seen a clear advantage of reducing convolutional feature maps such that central information is weighted more strongly. However, in the extreme case of completely removing everything but the center, we also see the prediction performances decline for early time points as well as for more central electrodes. This suggests that peripheral information is still necessary to explain a substantial amount of variance in the EEG data.

To test if spatial reweighting and then combining center and periphery information can further increase encoding performance, we compared the GCS model against the full and the center-crop model (see Fig. 2e). We found that the GCS model significantly outperforms the full model at time points later than 95 ms at occipital and temporal electrodes. For the earliest time point (95 ms), the full model slightly outperforms the GCS model at central electrodes. Interestingly, we found that the GCS model also outperforms the center-crop model at the later time points (119 and 144 ms) and thus, additionally improved predictions compared to the default, full model.

Overall, this suggests that a spatial reweighting can either outperform or perform on par with both the center-crop and the full model at almost all time points and electrodes, as well as outperform the periphery-crop model at all time points.

#### 3.1.4 Temporally distinct contributions of spatially distinct features

So far, we have shown that encoding models using either only center or peripheral information can outperform or perform on par with encoding models that use the full extent of the DNN visual feature maps. To better understand, whether spatially distinct regions of the DNN feature map explain the same or different variance in the EEG signal, we calculated the partial correlation for the predictions of the center-crop, periphery-crop, GCS, and full model, respectively, while using the predictions of one of the other models as the covariate (see Fig. 3a for a sketch of the methodology). Thus, variance that is explained by both model predictions is excluded from the correlation calculation. Partial correlations of center and periphery crops reveal distinct temporal patterns: the periphery-crop model explains more unique variance at early time points and less unique variance at time points between ∼120 and 145 ms (see Fig. 3b). In turn, the center-crop model explains unique variance delayed in time: the first significant time point is 93 ms for Iz and 102 ms for Pz compared to 76 ms and 75 ms for the periphery-crop model for Iz and Pz, respectively. The temporal delay of the center model is most prominent at occipital electrode Iz, while at the more central electrode Pz, the unique variance of the center-crop model with respect to the predictions of the periphery-crop model is substantially lower. These findings are in line with the previous findings shown in Fig. 2. However, using partial correlations we found direct evidence for unique temporal contributions of distinct spatial DNN features.

The GCS model explains unique variance over the center model at earlier time points as well as later time points at electrode Iz and Pz (see Fig. 3c). Furthermore, the GCS model explains unique variance over the periphery model at electrodes Iz and Pz at both early and late time points (see Fig. 3d). The periphery model also still explains some unique variance at very early time points, confirming again the link between peripheral information and early visual processing.

Lastly, we tested whether the unique variance explained by the GCS model over the center and periphery models, is indeed a contribution of the GCS-transform and is not present in the full model: thus, we tested the GCS model predictions against the full model predictions (see Fig. 3e) and find that the GCS model has substantially more unique variance than the full model at all time points for Iz. For Pz, the full model has more unique variance at very early time points, confirming yet again the results seen in Fig. 2e at 95 ms. Together, the results of the partial correlation analysis suggest that the processing of central and peripheral information is temporally distinct and that both temporal dynamics can be captured using retinal spatial reweighting.

### 3.2 Additional Experiment 1 - Center vs. Periphery

Using differential spatial sampling and spatial reweighting of DNN feature maps, we showed that central and peripheral information contribute uniquely to the explained variance of linearized encoding models predicting human EEG responses during natural scene viewing. To verify these distinct temporal profiles, we performed an additional experiment in which we explicitly stimulated center and periphery only (see Methods section 2.1.2). The experimental paradigm was kept fixed to a large extent with the main difference being that stimuli were presented using a circular aperture (that varied in diameter) that either excluded central information or excluded the directly inverse, peripheral information (see Fig. 4a for examples of stimuli).

We then built linearized encoding models to predict ERPs of held-out test images for data from each individual stimulus condition. Specifically, we tested: (I) whether we observe a temporal shift in the prediction performance of the central stimulation condition compared to the peripheral stimulation condition, and (II) whether the GCS feature transform can improve predictions even when only selectively stimulating central or peripheral regions. Thus, we built two encoding models for each stimulus condition, one encoding model used the full feature map (”Full”) whereas the other used the GCS-transformed feature maps (”GCS”).

#### 3.2.1 Experimental evidence for temporally distinct processing of center and periphery

In Fig. 4b), we report the corrected mean prediction performance across subjects, electrodes and time points later than stimulus onset. We find that on average, corrected prediction performances increase with increasing sizes of central stimulation, and decrease with increasing removal of central parts (periphery condition). Further, it appears as if the GCS model improves predictions compared to the full model for the small periphery condition and for the large center conditions. For the medium periphery condition, there seems to be a slight decrease in prediction performances while for the other conditions, the models appear to perform equally well.

Due to the limited number of subjects, we do not report group-level statistics but rather show the different model performances for each individual subject in Fig. 4c). Individual subject comparisons clearly show that performance is larger for the GCS model compared to the full model in the small periphery and large center condition in 3 out of 4 subjects.

Last, we show the mean corrected prediction performance across subjects for each stimulus condition and encoding model over time in Fig. 4d). We find a clear temporal shift between central and peripheral stimulation conditions for the full model. This confirms the previous observations from the modeling experiment that EEG signals recorded during peripheral stimulation can be explained well early in time, while signals obtained during central stimulation can be explained well later in time. When increasing the size of central stimulation, we observe an increase in overall encoding model performance for the full model but also a temporal shift towards earlier time points. Further, it appears as if the improvement of the GCS model for the small periphery condition is present at all time point after stimulus onset. For the large center condition, the GCS model seems to improve predictions at time points between 100 ms and 200 ms after stimulus onset compared to the full model.

Overall, we see temporally distinct profiles of processing central and peripheral information by selectively stimulating the regions of the visual field and find that good encoding performance requires sufficient stimulation of both central and peripheral regions.

### 3.3 Additional Experiment 2 - Stimulus Sizes

#### 3.3.1 Large-field stimulation reveals center-periphery temporal profiles

To test if spatial transformation of DNN feature maps aids encoding model performance in other datasets, we also ran a comparison of a standard (full) encoding model and the GCS encoding model on the publicly available EEG dataset by Gifford et al. (2022), containing responses for 10 subjects to 16,740 images from the THINGS dataset (Hebart et al., 2019). Unlike in our data, we see no improvement in encoding performance by including a GCS transformation at any electrode (see Supplementary Fig. S1). We hypothesised that this difference could reflect a lack of peripheral stimulation in the Gifford et al., dataset (image size 500×500 pixels, 7×7 degrees of visual angle), compared to our data (image size 2155×1440 pixels, 50×29.5 degrees of visual angle).

To test this hypothesis, we ran the second additional experiment (see Methods and Materials), whereby we systematically decreased the stimulus extent compared to the main experiment (small, medium, large; see Fig. 5a). Noise ceiling estimates of data from systematically varying stimulus extent show that the amount of signal in the data increases for increasing stimulus size (see Fig. 5b and Fig. S3 for ERPs for each conditions). Further, we find that there is indeed no benefit of GCS in the small condition, but a significant benefit for large stimuli (see Fig. 5c). Prediction performance for individual subjects show that the improvement of GCS over the full model is consistent across all four subjects for large stimulus sizes (see Fig. 5d). The results of this additional experiment suggests that the effects of retinal sampling on EEG responses are only revealed when using large-field stimulation that extends sufficiently into peripheral vision, thereby more closely approximating real-world vision.

## 4 Discussion

Building encoding models using DNN features allows for highly accurate predictions of neural signals during natural visual perception. However, linearly mapping the entire feature map onto the neural data assumes equal contribution of each spatial location when predicting the neural signal. Thus, the differential processing of central and peripheral input reflected in the structural and functional organization of human visual processing is not taken into account.

Here, we show that spatial selection of parts of the DNN feature maps improves encoding performance for predicting human EEG data. Using this approach, we unveiled distinct temporal profiles of peripheral vs. foveal visual information processing in the EEG signal. Next, we experimentally confirmed these distinct temporal dynamics of peripheral and foveal visual processing by selectively stimulating human vision at peripheral and foveal locations. Applying a GCS transform to the feature maps of the DNN improved encoding performances across both experiments and unified the predictivity of both early and late visual processing by spatially reweighting peripheral and central information. Last, we showed that the spatial and temporal dynamics underlying real-world visual processing can only be unveiled when approaching full-field visual stimulation.

### 4.1 Distinct temporal profiles of peripheral and foveal processing

The processing of the rich incoming visual information is guided by saccades that move the fovea onto locations that should be sampled with high acuity (Rosenholtz et al., 2012). Low-contrast and motion sensitive cells process peripheral information, thereby informing subsequent saccades while allowing for the detection of e.g., moving dangers. This idea is suggestive of a global to local processing theory.

A global to local processing hierarchy is thought to be implemented through the parvocellular and magnocellular pathways. The response properties of the parvo and magno cells have been studied extensively and have been shown to underlie the specialization of processing of foveal and peripheral information (Solomon, 2021). An important characteristic of the magnocellular pathway, which is largely processing peripheral information, is the processing of *scene gist* (Oliva, 2005). Scene gist refers to the idea of processing the context of a scene (Pereira and Castelhano, 2014) in very short time to facilitate self-orientation (Epstein, 2005; Malcolm et al., 2016), processing action affordances (Bonner and Epstein, 2017; Bartnik and Groen, 2023), and setting up priors for fast object recognition (Aldegheri et al., 2023).

Behavioral studies have shown the speed of scene gist processing (Oliva and Torralba, 2006) as well as the importance of peripheral stimulation for scene categorization tasks (Loschky et al., 2017). Further, EEG signals measured during RSVP experiments with humans have been shown to be highly predictable by image-computable scenes statistics (Groen et al., 2013). Recently, Jang and Tong (2024) showed that training DNNs on blurred images, thereby approximating the peripheral percept, improves both behavioral and representational alignment with humans.

Our findings are consistent with the idea of a fast, coarse processing of peripheral information followed by a delayed period during which the EEG signal is dominated by foveal stimulation, proposed in various global-to-local frameworks of visual processing (Hochstein and Ahissar, 2002; Bar, 2004; Hegdé, 2008). By spatially selecting parts of the DNN feature maps, we show that information at the center of a scene is predictive of time points later than 144 ms at occipital electrodes. This signal is likely dominated by the activity of neurons in the primary visual cortex that has receptive field properties that correspond to a fine, foveal information processing (Harvey and Dumoulin, 2011). By selecting parts of the DNN feature maps that correspond to peripheral scene information we show that this information is being processed at early time point and at more central and temporal electrodes. We provided additional experimental evidence that confirm these temporal profiles during selective stimulation of foveal and peripheral regions of the visual field.

### 4.2 Optimal spatial reweighting using retinal sampling

Next to work on modelling peripheral processing, many attempts have been made to incorporate foveation into computational models of visual processing. Behavioral benefits have been observed by foveating training images of DNNs (Deza and Konkle, 2020). Bringing together both foveal and peripheral spatial sampling, da Costa et al. (2024) showed that a spatial reweighting of DNN training images that magnifies foveal information yielded a hierarchy of receptive field properties that is similar to that of the human visual system. We find that the same spatial reweighting applied to pretrained DNN feature maps can improve encoding model performance thereby accounting for both early and late temporal dynamics in the EEG signal.

For certain electrodes, at central and temporal locations, we see that the periphery crop model explains unique variance that the GCS model does not capture. An important parameter in the GCS transform is the magnification factor of central regions. Performing hyperparameter optimization on this parameter for each electrode individually, and potentially even for each time point, could not only improve encoding model performance further, but could also yields new insights into the gradients of foveal over-representation in EEG recordings over time and across the scalp.

### 4.3 Studying real-world visual processing

Historically, visual information processing in the human brain has been investigated using hand-crafted stimuli that were shown to participants in carefully designed neuroimaging experiments, to derive response properties (Dumoulin and Wandell, 2008). However, applying these methods to real-world visual stimuli is often hard or not possible. With the advent of DNNs that are intrinsically image-computable, meaning they can process arbitrary natural images, a lot of attention has been put into studying visual processing in humans using DNNs and natural images. However, only few efforts (e.g., Park et al., 2023) have been made to extend this beyond the experimental setting of small stimulus sizes presented on low resolution screens during neuroimaging experiments. When using small size visual stimuli, only a small proportion of the visual field is stimulated. Stimulating only (para-)foveal regions does not require modeling of peripheral spatial sampling to accurately predict neural recordings. Here, we show that in scenarios where both foveal and peripheral regions are being stimulated however, the prominent differential sampling of foveal vs. peripheral information plays a significant role for accurately predicting neural data.

Moreover, we also show that exclusively stimulating foveal regions reduces the signal-to-noise-ratio (SNR) of the EEG recordings compared to large-field stimulation. One possible explanation for these differences in SNR is that the lack of activation in the proportion of cortical volume that processes peripheral information increases the noise in the EEG signal when peripheral stimulation is absent. Interestingly, our results suggest that the increased SNR in the EEG signal during large-field stimulation cannot be explained anymore by using the standard encoding model approach of mapping the entire feature map onto the neural data. Only when accounting for the differential spatial sampling of foveal and peripheral regions, is it possible to explain the additional explainable variance.

### 4.4 Limitations and suggestions for future work

The analysis of EEG recordings from the main experiment could have benefited from an increased number of repetitions per image condition as well as number of image conditions which have been shown to increase SNR (see e.g., Gifford et al., 2022). Recording eye-tracking data in the additional experiment could have allowed for exclusion of epoch during which participants did not maintain fixation at the center of the screen. Specifically, this could influence the SNR during periphery conditions where participants were asked to fixate in the center of the screen while stimuli were only presented at peripheral locations, thus potentially soliciting involuntary saccades. Further, the two additional experiments comprise a small number of subjects. Extending this to more subjects would allow for statistical tests of effects between encoding model performances across subjects.

The employed method of retinal spatial reweighting used here was originally introduced as a transform of the input to the DNN instead of to the DNN feature maps. The input-level transformation would be a more biologically plausible implementation, posing processing within the DNN as cortical processing and the GCS as a retinal (i.e., prior to the cortex) transform. We performed these experiments, i.e., we applied the GCS at the input level and found that encoding model performance would also increase compared to using non-transformed input (results not show here). However, testing task-performance (here, object recognition) using the GCS transform of input images, revealed that inputs appeared to be shifted out of distribution (OOD) compared to the training data, thus showing a substantial drop in DNN performance. Applying the GCS transform to DNN feature maps instead, allows for maintaining task-performance while also increasing encoding model performance of EEG signals.

An exciting way of resolving this problem would be to incorporate the GCS transform already during the task training of the DNN thus preventing a possible OOD shift during feature extraction for the encoding model. Our custom dataset of outdoor scenes, which additionally feature object bounding box annotations, would allow for training of the DNN on this dataset. Crucially, the GCS transform could be used to model not only foveal magnification, but also to simulate saccades by centering the GCS transform on the object instead of on the center of the scene. While these experiments are out of scope of the current paper, we will make the OADS image dataset publicly available upon publication, such that future work can explore these directions.

Overall, our results suggest that a fruitful approach of improving encoding models using DNN feature maps is the implementation of biological knowledge about the human visual system into the encoding model.

## 5 Acknowledgements

This work is supported by the Interdisciplinary PhD Programme of University of Amsterdam Data Science Center.

## 6 Contributions

N.M. performed analyses. N.M., H.S.S., and I.I.A.G. wrote the paper. N.M., H.S.S., and I.I.A.G.. originally conceived the approach.

## 7 Supplementary Information

**Figure S1:**
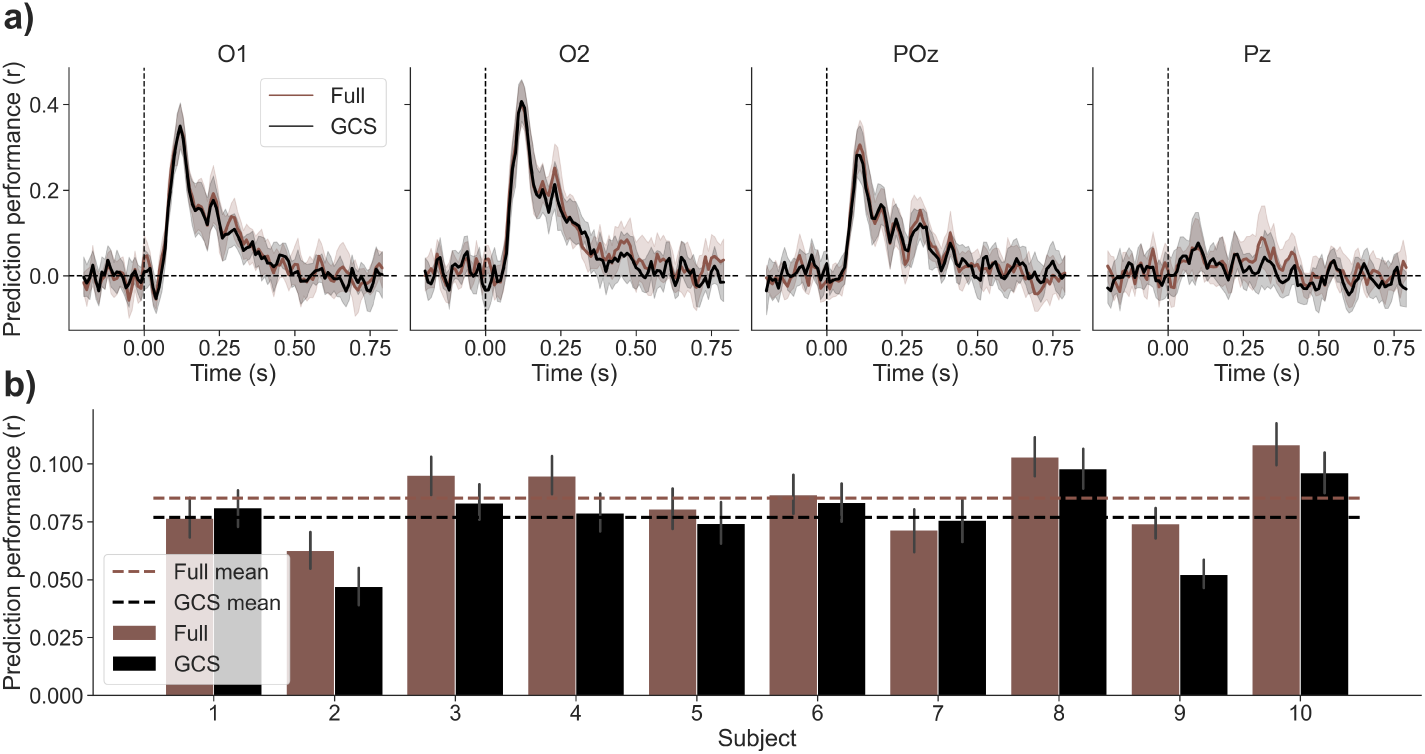
GCS model performs equally well as full model on large public EEG dataset (Gifford et al., 2022). **a)** Mean encoding model performance across subject for full (brown) and GCS (black) model over time for four representative electrodes over time. Shaded areas indicate 95% confidence interval per time point across subjects. At no time point for the selected electrodes does there appear to be an advantage of either model over the respective other. **b)** Mean encoding model performance across electrodes per subject for full (brown) and GCS (black) models averaged across all time point after stimulus onset. Dashed lines indicate subject average per encoding model. It seems as if no consistent difference between encoding model performances appear across subjects.

**Figure S2:**
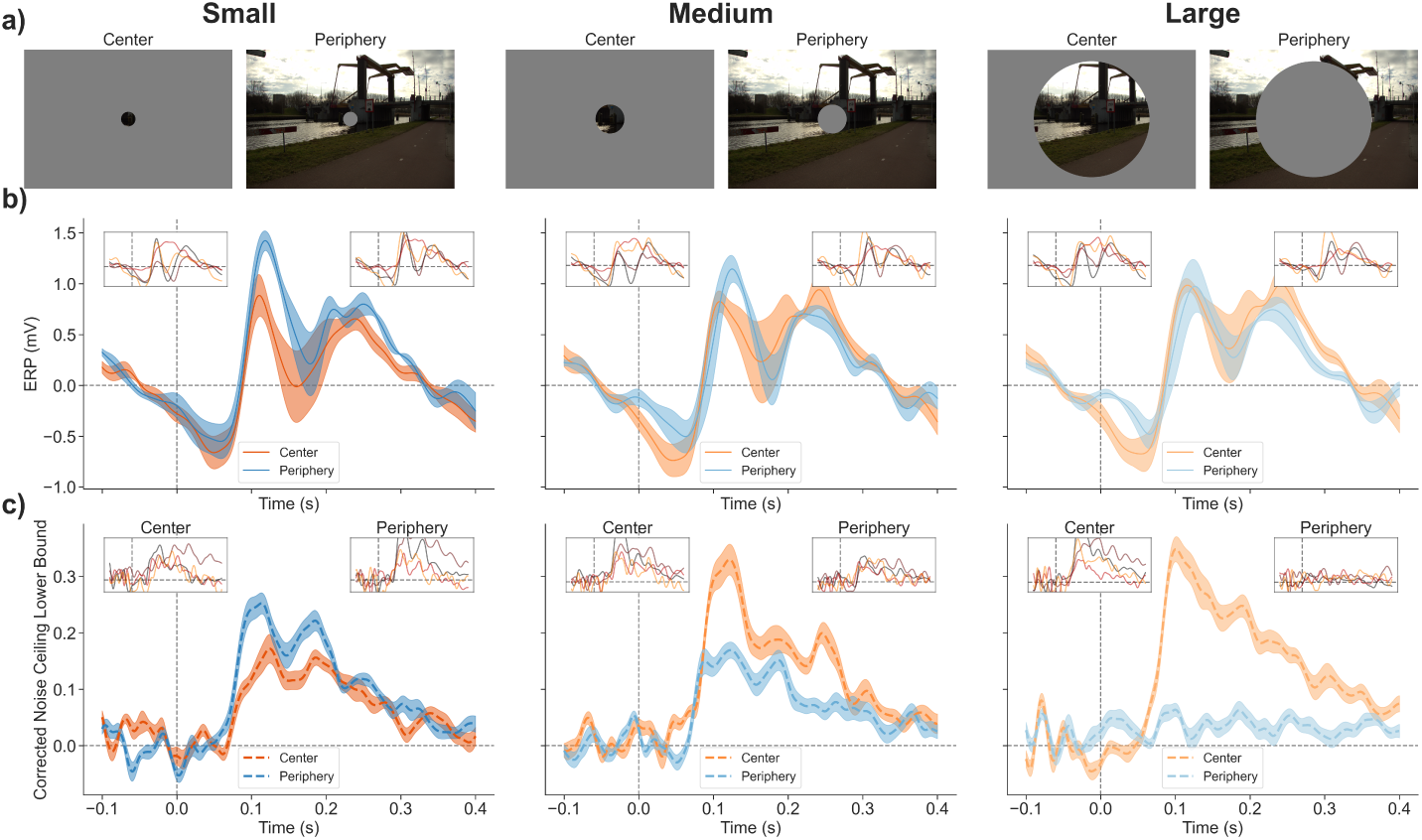
Distinct neural signatures of central vs. peripheral stimulation. **a)** Example stimuli for central vs. peripheral stimulation using a circular aperture increasing size, from left to right. **b)** Average ERPs across subjects for center (orange) and periphery (blue) conditions for increasing aperture sizes. Axis insets show per subject ERPs for center (top-left) and periphery (top-right). ERPs are higher for peripheral stimulation at time point between 100 and 150 ms after stimulus onset. **c)** Corrected noise ceiling lower bound estimate averaged across subjects for center (orange) and periphery (blue) conditions for increasing aperture sizes. Axis insets show per subject noise ceiling lower bound estimates for center (top-left) and periphery (top-right). For small apertures, peripheral stimulation yields a higher SNR earlier in time compared to central stimulation. With increasing aperture size, SNR for central stimulation increases while SNR for peripheral stimulation decreases.

**Figure S3:**
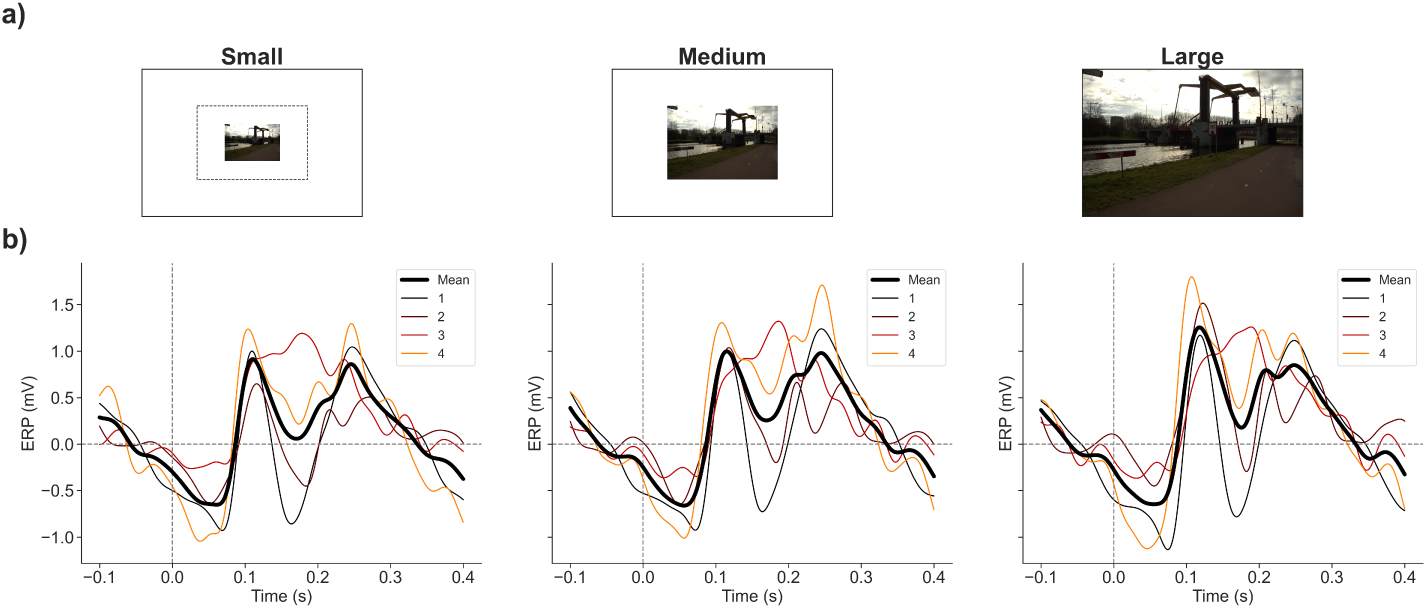
**a)** Example stimuli for small, medium, and large stimulus sizes, from left to right. **b)** Average ERPs across 16 posterior electrodes for each subject individually (colored lines) plus subject averaged (thick black line) for increasing stimulus size conditions.

## References

Aldegheri, G., Gayet, S., and Peelen, M. V. (2023). Scene context automatically drives predictions of object transformations. Cognition, 238:105521.

Bar, M. (2004). Visual objects in context. Nature Reviews Neuroscience, 5(8):617–629.

Bartnik, C. G. and Groen, I. I. (2023). Visual perception in the human brain: How the brain perceives and understands real-world scenes. In Oxford Research Encyclopedia of Neuroscience. Oxford University Press.

Benjamini, Y. and Hochberg, Y. (1995). Controlling the false discovery rate: a practical and powerful approach to multiple testing. Journal of the Royal statistical society: series B (Methodological), 57(1):289–300.

Biederman, I. (1981). Do background depth gradients facilitate object identification? Perception, 10(5):573–578.

Bonner, M. F. and Epstein, R. A. (2017). Coding of navigational affordances in the human visual system. Proceedings of the National Academy of Sciences, 114(18):4793–4798.

Burr, D. and Ross, J. (1986). Visual processing of motion. Trends in Neurosciences, 9:304–307.

Cadieu, C. F., Hong, H., Yamins, D. L., Pinto, N., Ardila, D., Solomon, E. A., Majaj, N. J., and DiCarlo, J. J. (2014). Deep neural networks rival the representation of primate it cortex for core visual object recognition. PLoS computational biology, 10(12):e1003963.

Conwell, C., Prince, J. S., Kay, K. N., Alvarez, G. A., and Konkle, T. (2022). What can 1.8 billion regressions tell us about the pressures shaping high-level visual representation in brains and machines? BioRxiv, pages 2022–03.

Cowey, A. and Rolls, E. (1974). Human cortical magnification factor and its relation to visual acuity. Experimental Brain Research, 21:447–454.

da Costa, D., Kornemann, L., Goebel, R., and Senden, M. (2024). Convolutional neural networks develop major organizational principles of early visual cortex when enhanced with retinal sampling. Scientific Reports, 14(1):8980.

Delorme, A., Richard, G., and Fabre-Thorpe, M. (1999). Rapid processing of complex natural scenes: A role for the magnocellular visual pathways? Neurocomputing, 26:663–670.

Deza, A. and Konkle, T. (2020). Emergent properties of foveated perceptual systems. arXiv preprint arXiv:2006.07991.

Doerig, A., Sommers, R. P., Seeliger, K., Richards, B., Ismael, J., Lindsay, G. W., Kording, K. P., Konkle, T., Van Gerven, M. A., Kriegeskorte, N., et al. (2023). The neuro-connectionist research programme. Nature Reviews Neuroscience, 24(7):431–450.

Dumoulin, S. O. and Wandell, B. A. (2008). Population receptive field estimates in human visual cortex. Neuroimage, 39(2):647–660.

Elmoznino, E. and Bonner, M. F. (2024). High-performing neural network models of visual cortex benefit from high latent dimensionality. PLOS Computational Biology, 20(1):e1011792.

Epstein, R. (2005). The cortical basis of visual scene processing. Visual Cognition, 12(6):954–978.

Gifford, A. T., Dwivedi, K., Roig, G., and Cichy, R. M. (2022). A large and rich eeg dataset for modeling human visual object recognition. NeuroImage, 264:119754.

Gramfort, A., Luessi, M., Larson, E., Engemann, D. A., Strohmeier, D., Brodbeck, C., Goj, R., Jas, M., Brooks, T., Parkkonen, L., et al. (2013). Meg and eeg data analysis with mne-python. Frontiers in neuroscience, page 267.

Gratton, G., Coles, M. G., and Donchin, E. (1983). A new method for off-line removal of ocular artifact. Electroencephalography and clinical neurophysiology, 55(4):468–484.

Groen, I. I., Ghebreab, S., Lamme, V. A., and Scholte, H. S. (2012). Spatially pooled contrast responses predict neural and perceptual similarity of naturalistic image categories. PLoS Computational Biology.

Groen, I. I., Ghebreab, S., Prins, H., Lamme, V. A., and Scholte, H. S. (2013). From image statistics to scene gist: evoked neural activity reveals transition from low-level natural image structure to scene category. Journal of Neuroscience, 33(48):18814– 18824.

Grootswagers, T., Robinson, A. K., and Carlson, T. A. (2019). The representational dynamics of visual objects in rapid serial visual processing streams. NeuroImage, 188:668–679.

Güçlü, U. and Van Gerven, M. A. (2015). Deep neural networks reveal a gradient in the complexity of neural representations across the ventral stream. Journal of Neuroscience, 35(27):10005–10014.

Harvey, B. M. and Dumoulin, S. O. (2011). The relationship between cortical magnification factor and population receptive field size in human visual cortex: constancies in cortical architecture. Journal of Neuroscience, 31(38):13604–13612.

Hebart, M. N., Dickter, A. H., Kidder, A., Kwok, W. Y., Corriveau, A., Van Wicklin, C., and Baker, C. I. (2019). Things: A database of 1,854 object concepts and more than 26,000 naturalistic object images. PloS one, 14(10):e0223792.

Hegdé, J. (2008). Time course of visual perception: coarse-to-fine processing and beyond. Progress in neurobiology, 84(4):405–439.

Hochstein, S. and Ahissar, M. (2002). View from the top: Hierarchies and reverse hierarchies in the visual system. Neuron, 36(5):791–804.

Holmes, D. L., Cohen, K. M., Haith, M. M., and Morrison, F. J. (1977). Peripheral visual processing. Perception & Psychophysics, 22(6):571–577.

Intraub, H. (1981). Rapid conceptual identification of sequentially presented pictures. Journal of Experimental Psychology: Human Perception and Performance, 7(3):604.

Jang, H. and Tong, F. (2024). Improved modeling of human vision by incorporating robustness to blur in convolutional neural networks. Nature Communications, 15(1):1989.

Kazemian, A., Elmoznino, E., and Bonner, M. F. (2024). Convolutional architectures are cortex-aligned de novo. bioRxiv, pages 2024–05.

Keysers, C., Xiao, D.-K., Földiák, P., and Perrett, D. I. (2001). The speed of sight. Journal of cognitive neuroscience, 13(1):90–101.

Krizhevsky, A., Sutskever, I., and Hinton, G. E. (2012). Imagenet classification with deep convolutional neural networks. Advances in neural information processing systems, 25.

Kwon, M. and Liu, R. (2019). Linkage between retinal ganglion cell density and the nonuniform spatial integration across the visual field. Proceedings of the National Academy of Sciences, 116(9):3827–3836.

Loschky, L. C., Nuthmann, A., Fortenbaugh, F. C., and Levi, D. M. (2017). Scene perception from central to peripheral vision. Journal of vision, 17(1):6–6.

Malcolm, G. L., Groen, I. I., and Baker, C. I. (2016). Making sense of real-world scenes. Trends in cognitive sciences, 20(11):843–856.

Oliva, A. (2005). Gist of the scene. In Neurobiology of attention, pages 251–256. Elsevier.

Oliva, A. and Torralba, A. (2006). Building the gist of a scene: The role of global image features in recognition. Progress in brain research, 155:23–36.

Oyster, C. W., Takahashi, E. S., and Hurst, D. C. (1981). Density, soma size, and regional distribution of rabbit retinal ganglion cells. Journal of Neuroscience, 1(12):1331–1346.

Park, J., Soucy, E., Segawa, J., Mair, R., and Konkle, T. (2023). Ultra-wide angle neuroimaging: insights into immersive scene representation. bioRxiv, pages 2023–05.

Paszke, A., Gross, S., Massa, F., Lerer, A., Bradbury, J., Chanan, G., Killeen, T., Lin, Z., Gimelshein, N., Antiga, L., et al. (2019). Pytorch: An imperative style, high-performance deep learning library. Advances in neural information processing systems, 32.

Peirce, J., Gray, J. R., Simpson, S., MacAskill, M., Höchenberger, R., Sogo, H., Kast-man, E., and Lindeløv, J. K. (2019). Psychopy2: Experiments in behavior made easy. Behavior research methods, 51:195–203.

Pereira, E. J. and Castelhano, M. S. (2014). Peripheral guidance in scenes: The interaction of scene context and object content. Journal of Experimental Psychology: Human Perception and Performance, 40(5):2056.

Perrin, F., Pernier, J., Bertrand, O., and Echallier, J. F. (1989). Spherical splines for scalp potential and current density mapping. Electroencephalography and clinical neurophysiology, 72(2):184–187.

Potter, M. C. (1976). Short-term conceptual memory for pictures. Journal of experimental psychology: human learning and memory, 2(5):509.

Rodieck, R. (1979). Visual pathways. Annual Review of neuroscience.

Rosenholtz, R. (2016). Capabilities and limitations of peripheral vision. Annual review of vision science, 2(1):437–457.

Rosenholtz, R., Huang, J., Raj, A., Balas, B. J., and Ilie, L. (2012). A summary statistic representation in peripheral vision explains visual search. Journal of vision, 12(4):14– 14.

Schyns, P. G. and Oliva, A. (1994). From blobs to boundary edges: Evidence for time-and spatial-scale-dependent scene recognition. Psychological science, 5(4):195– 200.

Seeliger, K., Fritsche, M., Güçlü, U., Schoenmakers, S., Schoffelen, J.-M., Bosch, S. E., and Van Gerven, M. (2018). Convolutional neural network-based encoding and decoding of visual object recognition in space and time. NeuroImage, 180:253–266.

Solomon, S. G. (2021). Retinal ganglion cells and the magnocellular, parvocellular, and koniocellular subcortical visual pathways from the eye to the brain. In *Handbook of Clinical Neurology*, volume 178, pages 31–50. Elsevier.

Storrs, K. R., Kietzmann, T. C., Walther, A., Mehrer, J., and Kriegeskorte, N. (2021). Diverse deep neural networks all predict human inferior temporal cortex well, after training and fitting. Journal of cognitive neuroscience, 33(10):2044–2064.

Sucholutsky, I., Muttenthaler, L., Weller, A., Peng, A., Bobu, A., Kim, B., Love, B. C., Grant, E., Achterberg, J., Tenenbaum, J. B., et al. (2023). Getting aligned on representational alignment. arXiv preprint arXiv:2310.13018.

Trouilloud, A., Kauffmann, L., Roux-Sibilon, A., Rossel, P., Boucart, M., Mermillod, M., and Peyrin, C. (2020). Rapid scene categorization: From coarse peripheral vision to fine central vision. Vision Research, 170:60–72.

Vallat, R. (2018). Pingouin: statistics in python. J. Open Source Softw., 3(31):1026.

van Gerven, M. A. (2017). A primer on encoding models in sensory neuroscience. Journal of Mathematical Psychology, 76:172–183.

Vater, C., Wolfe, B., and Rosenholtz, R. (2022). Peripheral vision in real-world tasks: A systematic review. Psychonomic bulletin & review, 29(5):1531–1557.

Virtanen, P., Gommers, R., Oliphant, T. E., Haberland, M., Reddy, T., Cournapeau, D., Burovski, E., Peterson, P., Weckesser, W., Bright, J., van der Walt, S. J., Brett, M., Wilson, J., Millman, K. J., Mayorov, N., Nelson, A. R. J., Jones, E., Kern, R., Larson, E., Carey, C. J., Polat, İ., Feng, Y., Moore, E. W., VanderPlas, J., Laxalde, D., Perktold, J., Cimrman, R., Henriksen, I., Quintero, E. A., Harris, C. R., Archibald, A. M., Ribeiro, A. H., Pedregosa, F., van Mulbregt, P., and SciPy 1.0 Contributors (2020). SciPy 1.0: Fundamental Algorithms for Scientific Computing in Python. Nature Methods, 17:261–272.

Wässle, H., Grünert, U., Röhrenbeck, J., and Boycott, B. B. (1989). Cortical magnification factor and the ganglion cell density of the primate retina. Nature, 341(6243):643– 646.

Yamins, D. L., Hong, H., Cadieu, C. F., Solomon, E. A., Seibert, D., and DiCarlo, J. J. (2014). Performance-optimized hierarchical models predict neural responses in higher visual cortex. Proceedings of the national academy of sciences, 111(23):8619–8624.

